# Genome-wide association mapping within a single *Arabidopsis thaliana* population reveals a richer genetic architecture for defensive metabolite diversity

**DOI:** 10.1101/2021.09.13.460136

**Authors:** Andrew D. Gloss, Amélie Vergnol, Timothy C. Morton, Peter J. Laurin, Fabrice Roux, Joy Bergelson

**Author notes:** Corresponding author: Joy Bergelson.

## Abstract

A paradoxical finding from genome-wide association studies (GWAS) in plants is that variation in metabolite profiles typically maps to a small number of loci, despite the complexity of underlying biosynthetic pathways. This discrepancy may partially arise from limitations presented by geographically diverse mapping panels. Properties of metabolic pathways that impede GWAS by diluting the additive effect of a causal variant, such as allelic and genic heterogeneity and epistasis, would be expected to increase in severity with the geographic range of the mapping panel. We hypothesized that a population from a single locality would reveal an expanded set of associated loci. We tested this in a French *Arabidopsis thaliana* population (< 1 km transect) by profiling and conducting GWAS for glucosinolates, a suite of defensive metabolites that have been studied in depth through functional and genetic mapping approaches. For two distinct classes of glucosinolates, we discovered more associations at biosynthetic loci than previous GWAS with continental-scale mapping panels. Candidate genes underlying novel associations were supported by concordance between their observed effects in the TOU-A population and previous functional genetic and biochemical characterization. Local populations complement geographically diverse mapping panels to reveal a more complete genetic architecture for metabolic traits.

## 1. Introduction

Plants produce a vast array of structurally diverse secondary metabolites that collectively underpin a variety of functions -- from regulating growth and development, to tolerating abiotic stresses, attracting pollinators, and deterring pathogens and herbivores [1]. Illuminating the genetic architecture of secondary metabolism is not only integral to understanding plant physiology, adaptation, and diversity across environments [2]; it also provides precise routes to breed or engineer more durable and productive crops [3].

In recent years, genome-wide association studies (GWAS) have emerged as a tool of choice for elucidating the genotype-to-phenotype links that shape plant metabolic diversity [3–5]. GWAS involve tests for statistical associations between genetic variants and organismal phenotypes. Because they require only genotypic and phenotypic information across a panel of natural plant genotypes (accessions), GWAS offer a straightforward and efficient method for inferring genotype-to-phenotype links from datasets of millions of SNPs across the genome and thousands of metabolites, enabled by the parallel advances in genome sequencing and metabolomic profiling.

A paradoxical pattern emerging from the application of GWAS to plant metabolic features, however, is that only a few loci are associated with variation in the abundance of a given metabolite [5]. Indeed, an average of fewer than two significant loci per metabolite were discovered across four GWAS studies encompassing >6,500 metabolites in leaves and/or seeds of *Arabidopsis*, rice, and maize (N = 305-529 plant accessions per study) [6–9]. Such simple genetic architectures are surprising given that secondary metabolites are often the product of biosynthetic pathways that have many enzyme-catalyzed steps, as well as the capacity to interact with additional pathways [10]. On the other hand, some physical and topological properties inherent to biosynthetic pathways predict that mutations in certain genes will have outsized effects, and thus impose evolutionary constraints unevenly across genes in a pathway [11,12]. While this heterogeneity may help explain the simple genetic architectures revealed for metabolites by GWAS, it’s also clear that some true signals are lost. In particular, GWAS fails to replicate many functionally-validated loci uncovered through other techniques for interrogating the genetic basis of metabolic variation, such as QTL mapping [13].

Much attention has been paid to forces that reduce the efficacy of GWAS, and to both experimental designs and statistical approaches to mitigate them [14,15]. One relatively understudied factor is the composition of the mapping panel, especially the geographic distribution over which accessions are drawn [14,15]. This is an important consideration because GWAS mapping panels in plants have conventionally been assembled over broad geographic scales, such as the *Arabidopsis* Regional Mapping Population (RegMap) and 1001 Genomes Project (1001G), which are composed predominantly of accessions collected across the European continent [16,17]. This design ensures that a broad swath of the species’ genetic diversity is included within the mapping panel, one of the main advantages of GWAS compared to QTL mapping. However, it also exposes analyses to a variety of geographically-driven confounding forces.

The most popularized cause of confounding driven by geography concerns population structure [18,19]. False positive associations arise at non-causal variants whose genotypes are correlated (i.e., in long-range linkage disequilibrium) with causal variants, and geographic population structure is a major source of these correlations [18]. Accounting for differences in relatedness among accessions (e.g., through the inclusion of a relatedness matrix in the GWAS model) controls these spurious associations [20,21], but at the cost of reducing power to detect causal variants whose geographic distribution tracks major axes of population structure [22,23]. This limitation is likely to be more prevalent for traits underlying local adaptation over broad geographic scales [15], which may make it particularly relevant for specialized metabolites.

However, even with effective control for the effects of long-range linkage disequilibrium, additional confounding factors are strengthened in geographically structured populations. Three processes in particular can dilute the strength of association at a causal variant. First, many alleles have geographically restricted distributions, causing the genetic basis of a trait to vary across regions (genetic heterogeneity) [14,24,25]. A variant’s phenotypic effect is thus diluted by averaging across these regions. Because rare alleles tend to be more geographically restricted [26], mapping within local or regional panels would have the benefit of elevating the frequencies of some rare alleles relative to their species-wide frequency, while eliminating others that are absent from the region. This would enhance the ability to detect rare, informative SNPs, at least in some regions. Second, a locus can have more than two functionally-distinct haplotypes (allelic heterogeneity), especially in geographically broad mapping panels that have high genetic diversity [14,27]. Because GWAS typically interrogates biallelic SNPs, a variant’s effect is diluted by averaging across the haplotypes tagged by each allele. Third, population structure across multiple causal loci can produce different genotypic combinations in different geographic regions. GWAS is less powerful when a causal variant’s effect is markedly weakened in some genetic backgrounds due to epistasis, since standard GWAS models are formulated to detect average additive effects across genetic backgrounds [28,29]. All of these factors point to the benefit of mapping in local panels, provided that adequate phenotypic and genetic variation is present.

Glucosinolates (GSLs), the primary class of secondary defensive metabolites in *Arabidopsis* and a model system for the genetics of plant secondary metabolism [30], offer a compelling opportunity to test the hypothesis that a local GWAS mapping population can better expose the genetic architecture of a complex trait than a geographically broad GWAS population. Glucosinolate biosynthesis has a polygenic basis, including a number of sequential enzyme-catalyzed reactions to produce a given aliphatic GSL (Methionine-derived, 12-15 reactions) or indolic GSL (Tryptophan-derived, 7-9 reactions) from their precursor amino acid [31]. Each step of the pathway has been functionally characterized through forward and reverse genetics approaches, leading to the identification of at least 45 genes involved (which is greater than the number of reactions due to functional redundancy among paralogs) [31]. Yet three GWAS of aliphatic GSL variation with large mapping populations (N > 300) spanning across Europe have consistently described associations at only three biosynthetic loci [6,13,32], even though the causal polymorphisms underlying mapped QTL have been localized to additional biosynthetic genes [33].

Intriguingly, conditions for all the sources of confounding detailed above are met for GSLs across the European distribution of *Arabidopsis [13]*. Recurrent loss of function and gene conversion events have generated complex patterns of allelic heterogeneity, including rare variants, and the geographically restricted distributions of functionally-defined haplotypes at a few major-effect loci implies strong genic heterogeneity [13,32,34]. Higher-order epistatic interactions among these loci determine which GSL molecules accumulate, resulting in GSL profiles that can be binned into qualitative “chemotypes,” defined by whether the gene(s) at each locus are functional [35]. Distributions of these epistatically-defined chemotypes are also geographically biased, displaying regional or continental clines shaped by a combination of demography and local adaptation [13,32]. If similar patterns have arisen at other loci with more modest phenotypic effects, geographic confounding might hinder their detection through GWAS; at the very least, large effect epistasis has been documented for other GSL biosynthetic enzymes [33,36]. Finally, even without geographic confounding, allelic heterogeneity, or epistasis, loss-of-function variants at major biosynthetic enzymes that are not captured well by polygenic genomic background effects in GWAS might add phenotypic noise that overwhelms modest signals of association at other loci.

Here, we quantified variation in GSL profiles in a single local population of *Arabidopsis*, compared the genetic architecture revealed through GWAS in this local population and in geographically broad mapping panels, and explored potential confounding factors underlying differences in the performance of the mapping populations. We focused on a population from Toulon-Sur-Arroux (TOU-A), France, which was collected along a fence line spanning only a few hundred meters [37]. Previous investigations found that the TOU-A population harbors less than 20% of the variants segregating at detectable frequencies in the 1001G, yet variants underlying heritable variation for a wide range of morphological, growth, defense, and fitness-related traits in TOU-A can be successfully mapped using GWAS in this local population [37,38]. We restricted our focus to genes with validated functions in GSL biosynthesis, broadly defined to include core structure formation, side-chain elongation, and secondary modification [31]. Decades of research has compiled a near-exhaustive catalog of the genes participating in these processes and their substrate specificities, providing functional data supporting novel associations that we uncovered at these loci. Overall, the expanded catalog of natural polymorphisms shaping GSL variation in the TOU-A population suggests that GWAS in local mapping populations could complement and expand the genetic architecture for metabolic variation revealed from geographically broad mapping panels.

## 2. Methods

### (a) Plant growth

To minimize maternal effects, seeds were harvested from 305 TOU-A accessions grown at 22°C with a 16:8h light:dark photoperiod, with 3wks vernalization at 4°C in 8h:16h light:dark to synchronize flowering, in fall 2017. For GSL profiling in mid-2019, seeds were sown on a 1:1 blend of nutrient retention (BM1) and seed germination (BM2) soil mixes (Berger, CA) in a complete randomized block design with four replicates of 294 accessions. After 4d stratification at 4°C, growth trays were moved to a chamber with white LED light (180-200 μmol·s-1) at 20°C in 10h:14h light:dark. Seedlings were thinned to one per cell 1wk after germination. Trays were rotated and bottom-watered every second day with fertilizer (15N-16P-17K) solution at 100 ppm N until harvesting at 21d.

### (b) GSL Extraction and Quantification

All liquid preparation and storage steps throughout the following protocol were conducted in polypropylene 96-well plates sealed with silicone cap mats. Entire rosettes were first clipped from the root, weighed, and directly submerged into 1.2 mL 80% methanol, which inhibits endogenous myrosinase activity [39]. After 2d dark incubation at ambient temperature, samples were centrifuged for 1m at 4000 × g, and the supernatant was transferred into a fresh plate and stored at -80°C. Immediately prior to GSL profiling, 240µL was evaporated with a 96- pin air drier in a fresh plate and redissolved in 120µL 25% methanol. This approach was chosen after favorable comparisons to alternative extraction methods with freezing and/or homogenization steps (see Supplemental Note).

GSL content was quantified with an Agilent 1200 Series HPLC machine coupled to an Agilent 6410 triple quadrupole mass spectrometer with parameters described in [40]. Samples were eluted with 0.1% formic acid in water (A) and 100% Acetonitrile (B) using the following separation gradient: 3.5 min of 99% A followed by a gradient from 99% to 65% A (1 to 35% B) over 12.5 min, and a wash with 99% B for 4 min with 5 min post-run re-equilibration to 99% A. The mass spectrometer was run in precursor negative-ion electrospray mode, monitoring all parent ions from m/z 350–520 with daughter ions of m/z 97, which correspond to the sulfate moiety of the GSL analytes. External standards (sinigrin, every 12th sample; and a GSL extract from a mixture of TOU-A genotypes, every 24th sample) interspersed throughout each run were monitored to ensure consistency. Individual GSLs were identified based on their fragmentation pattern and retention time [32] (Table S1). Intensities for each molecule were integrated using *MSnbase* v2.8.3 [41] and *xcms* v3.4.4 [42], using a customized approach that did not require delineating discrete peak boundaries and thus enabled increased sensitivity for low abundance molecules (see Supplemental Note).

### (c) Genotypes

Genotypes for the TOU-A population were obtained from [37]. Genotype data for the RegMap [16] and 1001G [17] datasets were obtained from [43]. For the 1001G dataset, this consisted of SNPs that were directly genotyped through whole-genome resequencing (WGS). For the RegMap panel, this consisted of SNPs that were directly genotyped with a 250K SNP chip and supported by WGS in resequenced accessions, and SNPs imputed by intersecting the RegMap chip genotypes and 1001G WGS genotypes. 2.8M SNPs with greater than 95% imputation accuracy were retained, which primarily excludes SNPs with low-frequency alleles.

### (d) Broad-Sense Heritability of GSLs

We fitted linear mixed models for log-transformed ion counts per milligram of leaf tissue using *lme4* [44], including random intercept effects for the accession identity and for the plate containing the sample during extraction and HPLC-MS/MS quantification. Heritability was estimated as the proportion of variance explained by accession identity after excluding variance explained by sample plate identity. Significance of accession identity was assessed by a likelihood ratio test with one degree of freedom. For published measurements of Regmap [32] and 1001G [13] accessions, an identical model was implemented using GSL abundances scaled by sample weights as reported by the authors.

### (e) GWA Mapping

To standardize comparisons across datasets, analyses were conducted identically for the TOU-A, 1001G, and RegMap datasets. First, best unbiased linear predictors (BLUPs) were extracted from the linear mixed models above; for one dataset [6] that pooled biological replicates, abundances from the single technical replicate per accession were used directly. Values were converted to z-scores so that GWAS would produce effect size estimates in units of phenotypic standard deviations. Second, GWAS were implemented as linear mixed models in GEMMA v0.98.1 [45], including a centered genetic relatedness matrix (-gk 1) to account for population structure. Significance per SNP was assessed by Wald Tests (-lmm 1).

Traits that were modeled separately for GWAS included (1) abundances of each of the heritable GSL molecules, and (2) log_2_-transformed ratios of the abundances of pairs of molecules with precursor:product relationships (Fig. S1). For indolic GSLs in TOU-A, we also implemented a multi-trait GWAS approach (multivariate linear mixed model, mvLMM [46]), which jointly models the relationships between the abundances of all detected molecules. Severe genomic inflation and/or algorithmic termination errors prevented the implementation of these models for other molecules and mapping panels. Unless otherwise stated, all GWAS excluded SNPs with minor allele frequency (maf) < 5% or missing genotypes in > 5% of the accessions (relaxed to 10% for TOU-A, which had more uncalled sites). We excluded a small number of GWAS exhibiting systematic genomic inflation as determined from the median *P*-value (λ > 1.04) or an excess of associated SNPs (98th percentile of genome-wide *P*-values < 0.01).

To search for significant associations harboring GSL biosynthetic loci, we used a recently compiled catalogue of functionally validated genes in the aliphatic and indolic GSL biosynthetic pathways ([31]; categories: side chain elongation, core structure synthesis, side chain modification). Because peaks of association at known GSL biosynthetic loci in previous GWAS reside tens or even hundreds of kb from the causal genes [13,32,34]--which may arise from extended causal haplotypes [34], structural variants, or intergenic regulatory variants--we defined candidate SNPs as those within 30kb of known biosynthetic genes. For the three loci with significant SNPs in our re-analysis of the 1001G and RegMap datasets, for which the causal genes are well-established, we further extended these windows in 10kb increments until they captured 90% of the SNPs within 0.5Mb of the known causal loci (AOP2/3, GS-OH, MAM1/3) that harbored significant associations with single GSL molecules or precursor:product ratios in those datasets.

### (f) Population Genetic Comparisons

Methods for all population genetic analyses are described in the Supplementary Methods.

## 3. Results

### (a) A deficit of rare alleles in the local TOU-A population

A population genetic comparison between TOU-A and the European 1001G accessions revealed favorable conditions for GWAS relative to geographically broad mapping panels. First, for the particular example of glucosinolates, we found that epistatic variation increases rapidly with geographical distance (Fig. 1a). Second, despite reduced overall diversity (1.9M SNPs in TOU-A vs. 11.5M SNPs in 1001G), the TOU-A population (1.3M) and 1001G panel (2.2M) had a relatively comparable number of common variants (defined here as biallelic SNPs with maf > 0.03). Indeed, a large fraction of common variants from the 1001G panel (2.2M) were also common in TOU-A (0.83M, 38%), indicating the reduced genetic diversity in TOU-A arises from a lessened contribution of rare variants. This was reflected in the allele frequency spectrum: after downsampling the 1001G to account for differences in sample size, the TOU-A population still displayed a less pronounced enrichment of rare relative to higher frequency variants (Figure 1b), resulting in higher genome-wide values of Tajima’s D (Fig. 1c). This strong reduction in rare variants is expected to reduce confounding effects of allelic heterogeneity in TOU-A, while the presence of many common variants suggests this does not come at the expense of drastically culling the polymorphisms that can be interrogated through GWAS.

**Figure 1.**
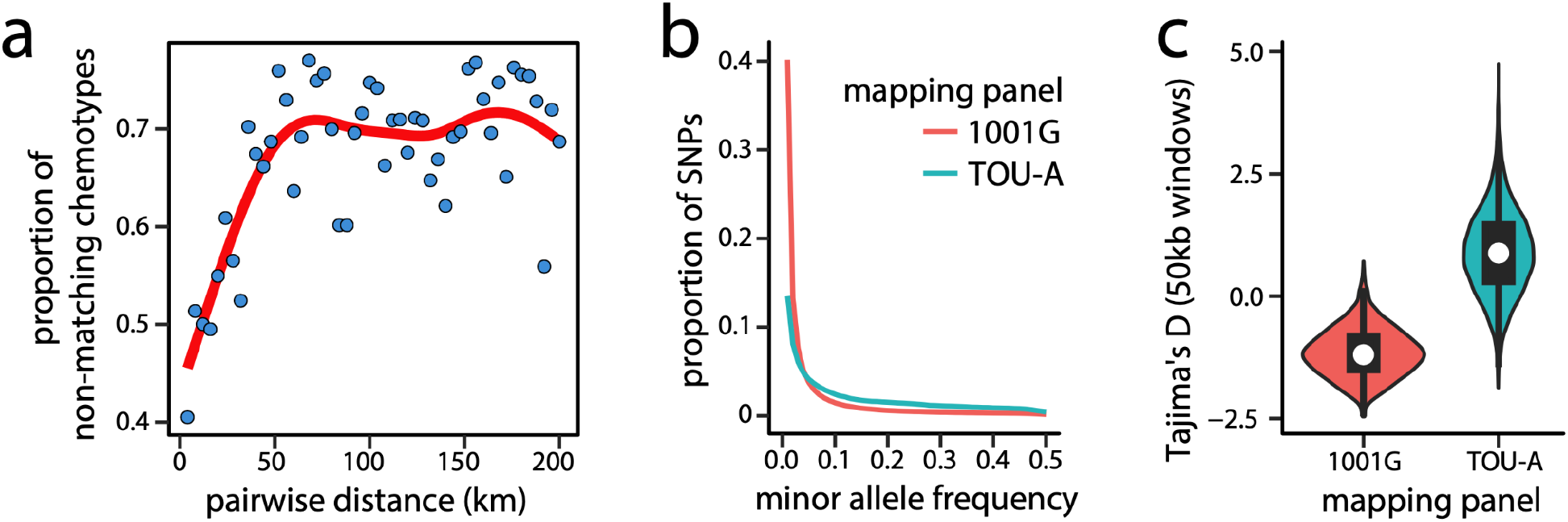
Reduced genetic complexity within local *Arabidopsis* populations. **(a)** The proportion of non- matching GSL chemotypes, which reflect the joint genotype at three epistatically-interacting loci (MAM, AOP, GS-OH), increases sharply and then plateaus as a function of geographic distance in pairwise comparisons among accessions. Points represent comparisons among European 1001G accessions in 4km bins. **(b)** The allele frequency spectrum is skewed toward common alleles in TOU-A relative to European accessions in the 1001G. The plotted lines were produced by connecting points indicating the proportion of SNPs falling into 1% bins of minor allele frequency. **(c)** Tajima’s D is also elevated in TOU-A, shown as a distribution of values across 50kb genomic windows. The 1001G panel was downsampled to 192 individuals to match TOU-A, and both populations were downsampled to 100 individuals per site, to avoid sample size and genotyping efficiency biases in panels b-c.

### (b) Heritable variation in glucosinolate profiles within the local TOU-A population

We quantified the relative concentrations of 13 major aliphatic and four indolic glucosinolates in 294 accessions from the TOU-A population under controlled growth chamber conditions. In contrast to broader geographic scales, where loss-of-function mutations within the glucosinolate biosynthetic pathway are pervasive, every TOU-A accession exhibited a fully functional GSL biosynthetic pathway. This was evidenced by abundant concentrations of the final products in the biosynthetic pathways for both short-chain aliphatic and indolic GSLs (Fig. S2).

Genetic differences among individuals explained significant portions of the between- accession variation in abundance for every GSL molecule: broad-sense heritabilities ranged from 0.19 < *H*^*2*^ < 0.92 (all *P*_*Bonferroni*_ < 0.05). In fact, analysis of GSL measurements from previous studies revealed systematically higher heritability estimates in TOU-A than the RegMap (Sign Test, median difference = 0.16 [95%CI:0.04,0.31], *P* = 0.02) and no significant difference between TOU-A and the 1001G (median difference = 0.04 [-0.20,0.20], *P* = 0.46) (Fig. 2). Although experimental design, tissue sampling, or data collection variables across studies could contribute to differences in heritability among the mapping populations, these data clearly indicate a high level of heritability for GSL traits within the local TOU-A population, even in the absence of the loss-of-function alleles at biosynthetic loci that have dramatic effects on GSL profiles across broader geographic scales.

**Figure 2.**
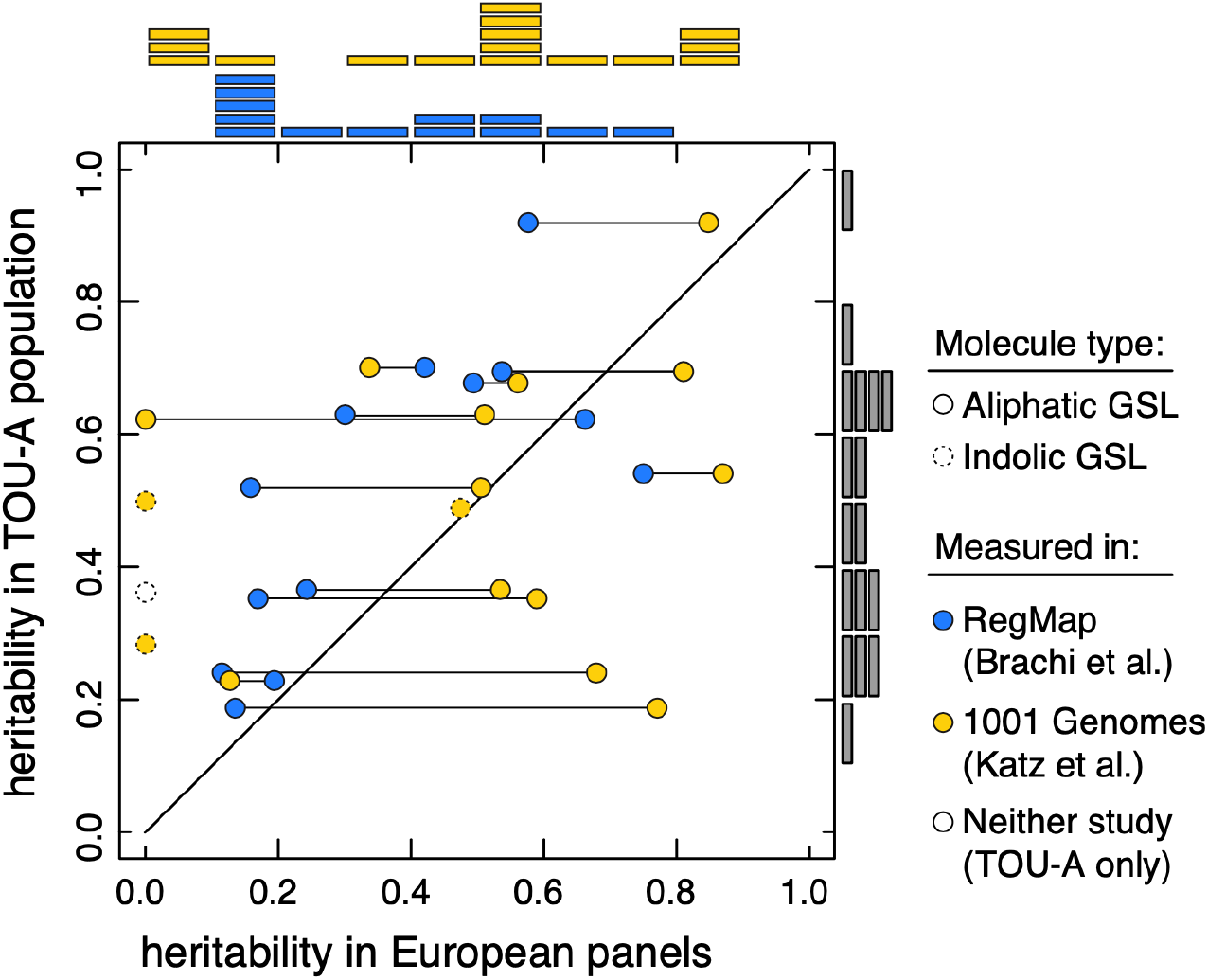
Glucosinolate variation is highly heritable within the TOU-A local population. **(a)** Estimates of broad-sense heritability (*H*^*2*^) for each GSL molecule in the TOU-A population are plotted against estimates in broader European mapping panels. Connected points indicate estimates of *H*^*2*^ for the same molecule in different European panels. Points above the diagonal line exhibit higher *H*^*2*^ in TOU-A. Histograms above and to the right of the plot indicate the distribution of *H*^*2*^ values in each population.

### (c) GWAS within the local TOU-A population reveals known and novel variants shaping aliphatic glucosinolate profiles

For 192 phenotyped accessions with whole genome sequences, we conducted GWAS using mixed models that controlled for confounding due to population structure by including a matrix of kinship among accessions as a random effect. We first focused on the abundances and relationships between 13 aliphatic GSLs.

#### Significant associations

The identity of associated loci in TOU-A depended on how GSL phenotypes were represented. Separate GWAS for the abundance of each molecule cumulatively uncovered significant associations at five biosynthetic loci (Fig. 3a). Given the strong positive and negative genetic correlations among GSL molecules in the TOU-A population (Fig. S3), we reasoned that mapping approaches utilizing these additional relationships may reveal additional associations. Indeed, using ratios of the abundances of individual precursor vs. product GSLs as the mapped traits cumulatively revealed significant associations at five biosynthetic loci, including two loci not recovered from GWAS using individual GSL abundances (Fig. 3b).

**Figure 3.**
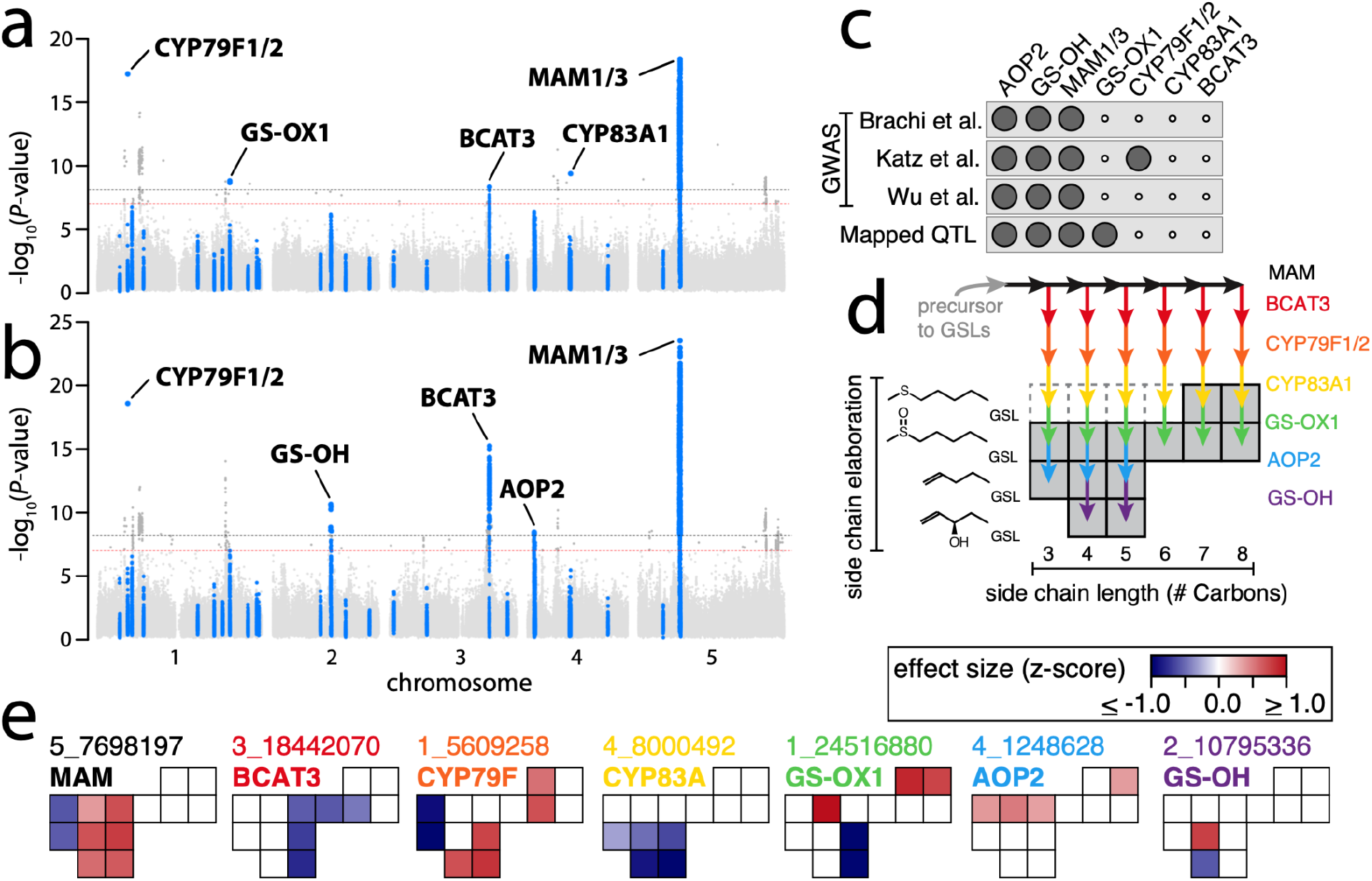
Seven biosynthetic loci are associated with aliphatic glucosinolate variation in the TOU-A local population. **(a**,**b)** The best *P*-value per SNP across individual GWAS, mapping either the abundance of individual GSL molecules (panel a, 13 traits) or the ratio of individual precursor vs. product molecule abundances (panel b, 17 traits). SNPs assigned to known GSL biosynthetic loci (see Methods) are enlarged and colored blue. Dotted lines indicate the Bonferroni genome-wide significance threshold for a single GWAS (red) or the full study (i.e., all individual GWAS across which *P-*values were merged; black). **(c)** For each locus associated with GSL variation in TOU-A, black circles indicate if the same locus was significant in GWAS in our re-analysis of GSL datasets from large (N > 300) European mapping populations [6,13,32] or was previously mapped as a QTL using biparental RILs [33]. **(d)** A model for how these loci interact to generate variation in GSL profiles for the major aliphatic GSLs present in TOU-A plants (shaded boxes). Enzyme-catalyzed reactions from precursor to product are shown as colored arrows. Dashed boxes indicate known intermediates that were not observed or quantifiable in TOU-A. **(e)** Effects on individual aliphatic GSLs for the minor allele of the leading SNP at each locus (identified as the SNP with the top association across any individual GWAS from panels a-b, named as “chromosome_position”). Boxes are oriented to represent the GSL molecules in panel d. Effect sizes are shown for each single molecule GWAS with *P* < 0.01 for the focal SNP.

The significant associations included the three loci (GS-OH, AOP, MAM) that we also recovered using the same approaches in a re-analysis of previous GWAS datasets, which consisted of mapping populations spanning the European continent (N > 300 accessions) (Fig. 3c & S4a). Many of these same associations were reported in the authors’ original analyses [6,13,32]. However, the GS-OX locus had not been mapped in the three GWAS with large mapping populations (although it was successfully mapped in biparental RILs) [33,47,48], and effects of natural polymorphisms in the BCAT3, CYP79F1, and CYP83A1 genes had not been described in any mapping study.

#### Effects on GSL profiles

A model for how the putatively causal enzymes at the seven significant loci generate GSL profile variation in the TOU-A population emerges simply by overlaying the reaction catalyzed by each enzyme, from precursor to product molecules, onto a plot of the major aliphatic GSLs detected in TOU-A plants. This produces a visual map of the variable steps in the biosynthetic pathway (Fig. 3d). We sought to use these relationships, supplemented with GSL profiles from gene knock-out mutants in previous studies, to validate each locus by comparing them to the effects inferred in our GWAS. To do this, we identified the leading SNP (i.e., the SNP with the strongest experiment-wide *P*-value) at each locus, extracted its GWAS model-fitted effect on the abundance of each GSL molecule, and visualized the effects on the map of GSL molecular variation in TOU-A (Fig. 3e). In addition to offering further evidence supporting the hypothesized causal genes at each locus, this approach illuminates how these loci generate different aspects of GSL profile variation in the TOU-A population.

The effects of the BCAT3 locus in TOU-A suggest that this gene underlies a dimension of variation in GSL side-chain length previously undescribed in natural populations of *Arabidopsis*, distinct from effects of the well-characterized variation at the MAM locus. The BCAT3 locus affected the abundances of GSLs with intermediate-length side chains, mirroring effects previously observed in a BCAT3 knockout mutant (Fig. 3e & S5). By contrast, functional genetic and biochemical assays have shown that the MAM1 and MAM2 enzymes primarily affect the abundance of GSLs with short side chains [49], similar to the inferred effect of the MAM locus in TOU-A, and MAM3 primarily affects the abundance of GSLs with long side chains (Fig. 3e & S5).

Of two previously unreported associations at cytochrome P450 monooxygenases functioning downstream of MAM and BCAT3 in the biosynthetic pathway (Fig. 3d), the novel association at the paralogous CYP79F1 and CYP79F2 genes [50] is especially noteworthy. The leading SNP at this locus was associated with a larger magnitude of effect on some short-chain molecules in TOU-A than MAM or BCAT3 (Fig. 3e), with especially large effects on molecules with the shortest observed side-chain length. This is consistent with the finding that among all biosynthetic enzymes, CYP79F2 exerts the strongest effect on pathway flux, with an outsized effect on propyl GSLs (i.e., GSLs with 3C side-chain lengths) [12]. Functional polymorphism at a CYP79F gene also underlies a QTL affecting the propyl fraction of GSLs in *Brassica juncea* [51], and separately underlies adaptive variation in the proportion of GSLs derived from branched-chain amino acids relative to methionine in *Boechera stricta [52]*. The association at CYP79F paralogs was recovered in our re-analysis of one European *Arabidopsis* dataset (Fig. S4), strengthening the evidence that CYP79F is a broadly important determinant of GSL profile variation across populations and species.

Two distinct loci harbor paralogous GS-OX genes that catalyze the *S*-oxygenation of methylthioalkyl to methylsulfinylalkyl GSLs with broad substrate specificity. While natural variation in the locus containing GS-OX2, GS-OX3, and GS-OX4 had been detected through QTL mapping with biparental RILs [47,48], neither locus had been detected in the three large, European GWAS panels. In addition to harboring a significant association when considering common variants (minor allele frequency, maf > 0.05; Fig. 3a), GS-OX1 harbored the strongest genome-wide association for many molecules when slightly rarer variants were considered (maf > 0.03; Fig. S6). Although biases in our GWAS model can yield inflated or deflated signals of association for alleles below this threshold, the strength of the association for this variant is exceptional even among alleles of similar frequency (0.05 > maf > 0.03). Intriguingly, the strongest associations at GS-OX1 did not involve methylthioalkyl GSL abundances individually or as a ratio compared to their derived methylsulfinylalkyl GSLs (Fig. S6), suggesting that linkage disequilibrium with other loci (or an unexpected effect of GS-OX1) may contribute to this association. Nevertheless, the effect on its direct precursor and/or product molecules is sufficient to drive a significant association: we further performed GWAS for a principal component capturing opposing shifts in the abundance of long-chain methylthioalkyl vs. methylsulfinylalkyl GSLs, and GS-OX1 harbored the strongest, statistically significant genome-wide association (Fig. S6).

Finally, effects of the two remaining polymorphisms in TOU-A, at the AOP [53] and GS-OH [54] loci, differed from the effects of loss-of-function variants at these loci that segregate over broad geographic scales, which eliminate the production of their GSL products and generate qualitative presence/absence variation in GSL profiles [13]. In TOU-A, by contrast, both loci affected their precursor GSL abundances, with only GS-OH also oppositely affecting (but not abolishing) its product GSL abundances (Fig. 3e).

It is important to note that the predicted effects do not include epistatic interactions, and that more subtle effects may not be discovered through GWAS. Accordingly, the effects described above should be interpreted only as the strongest, additive effects of each locus.

### (d) GWAS within the local TOU-A population reveals known and novel variants shaping indolic glucosinolate profiles

#### Significant associations

We implemented the same association mapping approach for four indolic GSL molecules, and were most successful when mapping traits that captured the relationships among abundances of different molecules. Three biosynthetic loci were significant in a multi-trait GWAS jointly modeling the abundance of all four indolic GSLs detected in TOU-A (Fig. 4a).

**Figure 4.**
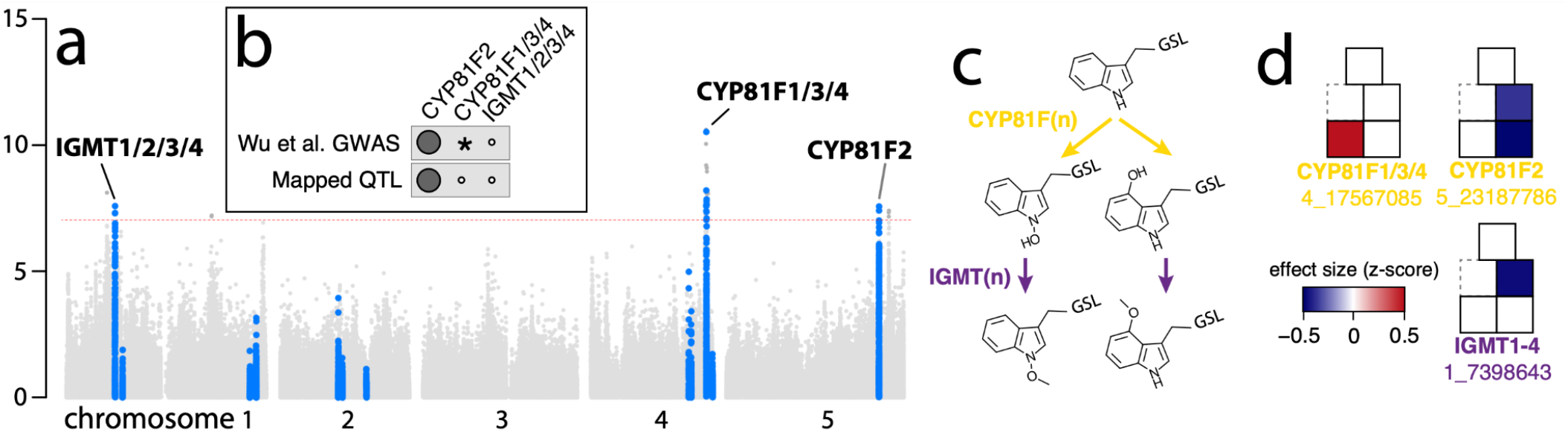
Three biosynthetic loci are associated with indolic glucosinolate variation in the TOU-A local population. **(a)** *P-*values from a multi-trait GWAS (mvLMM) jointly modeling all indolic GSL abundances. The plot layout, colors, and significance thresholds are as described in Figure 3a. **(b)** For each locus associated with GSL variation in TOU-A, black circles indicate if the same locus was significant in GWAS in our re-analysis of a GSL dataset from a large (N > 300) European mapping population [6] or was previously mapped as a QTL using biparental RILs [56]. “*” indicates a significant association in a published analysis that was not recovered in our standardized re-analysis. **(c)** The pathway for secondary modification of indole-3-ylmethyl GSL (top) through 1- or 4-hydroxylation (middle) and subsequent methoxylation (bottom). **(d)** Effects on individual indolic GSLs for the minor allele of the leading SNP at each locus, determined as in Fig. 3e. Boxes are oriented to represent the GSL molecules in panel c.

Of these three loci, two (both CYP81F loci) have been previously identified in GWAS [6] and remained the only two significant associations in our re-analysis of other datasets (Fig. 4b & S4). One of these loci was also discovered through QTL mapping, and CYP81F2 was functionally validated as the causal gene [55,56]. The IGMT locus had not been linked to natural variation in GSL profiles previously.

#### Effects on GSL profiles

Each putatively causal biosynthetic enzyme underlying the associations with indolic GSL variation in TOU-A has been functionally characterized through biochemical assays and in gene knockout mutants. CYP81F paralogs collectively catalyze the first elaboration step at different sites of indolic GSL ring structure [55,56], and IGMT paralogs collectively catalyze a subsequent elaboration step [57] (Fig. 4c). Using the effects of each locus extracted from our GWAS models, we looked for concordance between our GWAS (Fig. 4d) and previous QTL mapping, functional genetic, and knockout mutant studies to inform how these loci shape GSL variation in TOU-A.

The CYP81F subfamily of cytochrome P450 monooxygenases are responsible for hydroxylation of indolyl-3-ylmethyl (I3M) GSL [55,56], which can subsequently be methoxylated by other enzymes. The locus harboring CYP81F2 affected two GSL molecules in TOU-A (4-hydroxy-I3M-GSL and its derivative, 4-methoxy-I3M-GSL), which also differentially accumulate due to the CYP81F2 locus in a previous QTL mapping experiment [56]. The locus harboring CYP81F1, CYP81F3, and CYP81F4 paralogs affected the GSL that is methoxylated at a different site, 1-methoxy-I3M-GSL; the CYP81F-catalyzed product from which it derives, 1- hydroxy-I3M-GSL, is unstable and was not observable through our GSL profiling approach. These results further support evidence from previous mapping studies that paralogs at the two CYP81F loci affect different GSL molecules *in planta*, despite overlap in substrate specificities *in vitro* [55,56].

Four of the five indole glucosinolate O-methyltransferases (IGMT1-4) in *Arabidopsis* form a tandem array at the locus identified in our GWAS [57]. This locus had a strong effect on the abundance of its substrate, 4-hydroxy-I3M-GSL (Fig. 4d). Although IGMT1-4 enzymes cumulatively can methoxylate both 1- and 4-hydroxy-I3M-GSL in biochemical assays, our observation of effects restricted to 4-hydroxy-I3M-GSL methoxylation support a model previously inferred from the characterization of an IGMT5 knockout mutant, which retained functional copies of all four IGMT1-4 paralogs [57]. The mutant exhibited an absence of 1- methoxy-I3M-GSL but no reduction in 4-methoxy-I3M-GSL, suggesting the IGMT1-4 locus is responsible only for 4-methoxy-I3M-GSL’s production *in planta*.

Taken together, our results more fully link the functional variation characterized in enzyme biochemical and gene knockout studies with the variation for indolic GSLs observed in natural populations, identifying loci acting at three of the four secondary modification steps that give rise to the major I3M-derived GSLs in the TOU-A population.

### (e) Reduced population structure is unlikely to underlie improved performance of GWAS for glucosinolate profiles in the local TOU-A population

GSL profiles, and some of the large effect loci that underlie them, show strong geographic clines within and across Europe [13,32]. This raises the possibility that methods to control for population structure in GWAS could weaken signals of association with GSLs at loci whose genotypes are strongly correlated with population structure. To investigate this, we used ADMIXTURE to infer subgroups (k = 5) contributing to population structure separately within the TOU-A and the 1001G accessions. Focusing on the ten glucosinolate biosynthetic loci recovered by GWAS in TOU-A, we found that among-group variation in allele frequency was not elevated in the 1001G relative to TOU-A (Fig. S7). This suggests that the efficacy of GWAS for GSLs in TOU-A is unlikely to be the product of weaker population structure at causal loci, and may instead arise from differences in other confounding factors that are exaggerated in geographically broad mapping panels.

## 4. Discussion

As one of the best-studied secondary metabolite pathways in plants--with a wealth of functional genetic knowledge from GWAS and QTL mapping of natural variation, characterization of genetic mutant lines, and enzyme biochemical assays [30]--GSLs offered a compelling opportunity to investigate the performance of GWAS using a local mapping population. The expanded genetic architecture revealed for GSLs in the TOU-A population highlights the benefits of this approach. A modest mapping panel (N=192 accessions) led not only to the discovery of variants that were absent in geographically broad mapping panels with 1.5-4x more accessions, but also to novel loci whose contribution to natural variation was unknown despite numerous QTL mapping studies previously conducted for GSLs. These associations spanned each major portion of the pathway (Fig. 5): the MAM-catalyzed reaction loop for side-chain elongation in GSL precursor molecules, sequential steps for synthesis of the GSL core structure, and every level of secondary modification subsequent to the formation of a functional GSL molecule [31]. Thus, GWAS within a single population can offer a deep catalog of functional polymorphism within a biosynthetic pathway.

**Figure 5.**
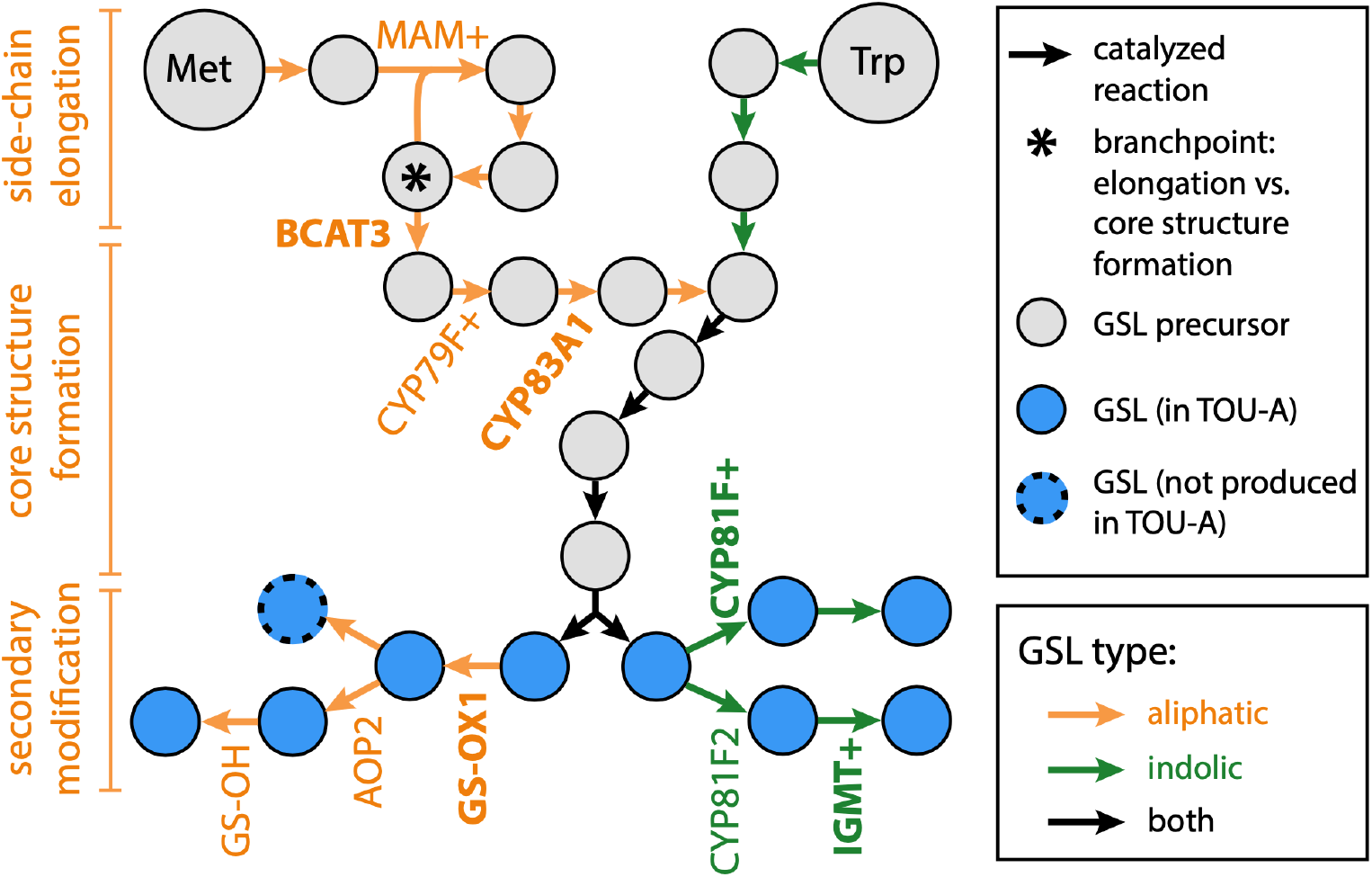
An overview of glucosinolate biosynthetic loci associated with GSL variation in the TOU-A population. The diagram shows each enzyme-catalyzed step, beginning with the amino acid precursor (Met or Trp). Genes harboring significant GWAS associations in TOU-A are listed at the biosynthetic step they catalyze. Bolded genes are novel associations, defined as those significantly associated in TOU-A but not in our re-analysis of three datasets with geographically broad European mapping panels. A “+” indicates that multiple paralogous genes at a locus could contribute to the association (e.g., CYP79F1 and CYP79F2 are represented as CYP79F+). The pathway and enzyme positions are based on [31]. Note that additional steps producing GSLs that accumulate only at very low levels in leaves are omitted.

The simplest explanation for the effectiveness of GWAS in TOU-A may be the observed reduction in genetic diversity relative to the broader European population. Theory predicts that allelic heterogeneity, which poses a major obstacle for GWAS, will be more pervasive in more genetically diverse populations. Further, the fact that diversity was reduced in TOU-A primarily through a relative deficit of rare variants, as expected if rare variants are geographically restricted and therefore locally more common [26], likely provides an additional benefit. Rare variants are not only poorly detected through GWAS, but their presence can obscure true associations at causal loci [58]. Consistent with this, GWAS has uncovered more associations and a broader (albeit largely unvalidated) functional repertoire of underlying candidate genes-- including biosynthetic enzymes, transcription factors, and transporters--across cultivars of *Brassica napus* than in European panels of *Arabidopsis* [59–61]. *B. napus* cultivars are less genetically diverse and have an excess of common variants (reflected in elevated Tajima’s D) relative to *Arabidopsis* [17,61,62], which may have been further exaggerated at glucosinolate-related genes by the diversity-reducing effects of directional selection during the breeding process [62].

While the general benefits of reduced geography-driven confounding in local populations should extend to GWAS for a variety of traits, our findings also illustrate properties of local populations likely to be especially beneficial when studying metabolite diversity specifically. In particular, the confounding effects of loss-of-function polymorphisms were absent from the major loci (MAM, AOP, GS-OH) that segregate such mutations over broad geographic scales. Loss-of-function mutations produce a particularly severe form of allelic heterogeneity. Many different mutations can produce analogous loss-of-function alleles at a gene, resulting in a high gene-wide mutation rate, such that many loss-of-function polymorphisms involve multiple haplotypes with parallel loss-of-function mutations [27]. Furthermore, loss-of-function mutations underlie dramatic epistatic effects, which may dilute additive effects modeled by GWAS. An extreme example involves the GS-OH locus that catalyzes the final secondary modification in the biosynthetic pathway (Fig. 5): loss of function alleles at upstream enzymes fully mask the effect of GS-OH on GSL variation in the majority of genetic backgrounds in *Arabidopsis*, and GS-OH itself segregates numerous loss-of-function alleles [13]. Of the three major large-effect loci mapped in other GWAS of aliphatic GSLs, only GS-OH has failed to consistently yield associations across previous analyses [6,13,32,34].

Although statistical approaches exist to mitigate geographically-driven confounding factors, they cannot entirely control for them. For example, GWAS models can be extended to include epistatic interactions alongside, or instead of, additive effects [63]. However, the immense number of possible pairwise interactions across the genome creates computational challenges and a severe multiple testing burden [64]. Other confounding factors can be lessened by altering genotype information rather than the GWAS models themselves. One simple yet powerful approach involves collapsing all predicted loss-of-function variants at a gene into a single allele, reducing their contribution to allelic heterogeneity [65]. Nevertheless, this approach requires genotyping to be conducted through whole-genome sequencing, and even then, many cases of abolished or altered gene function are difficult to annotate from DNA sequence data alone. Furthermore, while this approach can improve power to discover associations at loci with heterogeneous loss-of-function variants, it does not address their confounding epistatic effects on other loci. Even in cases where various genotyping and statistical approaches do largely succeed in mitigating specific confounding factors, integrating them to address many factors simultaneously is challenging. For many research questions, the use of local mapping populations in which these confounding factors are lessened offers an attractive alternative to these more tailored GWAS approaches.

Despite their benefits, GWAS in local populations are certainly not ideal for every research question. GWAS of GSLs in different mapping populations illustrate this clearly: integrating population genomic analyses with GWAS using *Arabidopsis* accessions sampled throughout Europe revealed how GSL profiles have been shaped by adaptation and demography across the species range [13,32,34], which would be impossible to infer from a single local population. Meanwhile, GWAS using the TOU-A population implicated more loci in natural phenotypic variation than could be detected in broader mapping panels. Complementary GWAS in local and geographically broad mapping panels thus provide an exciting avenue toward a fuller understanding of the genetic variation and evolutionary processes that shape phenotypic diversity in nature.

## 5. Data Accessibility

Raw data are accessible on the Dryad Digital Repository (https://doi.org/10.5061/dryad.4mw6m90b6). Scripts are available on GitHub (https://github.com/peterlaurin/TOUA_Glucosinolate_GWAS).

## 6. Authors’ Contributions

A.D.G., F.R., and J.B. conceived of the study. A.D.G, A.V., T.C.M., and J.B. collected the data. A.D.G, A.V., T.C.M., and P.J.L. analyzed the data. A.D.G. and J.B. wrote the manuscript.

## 8. Acknowledgments

We thank Xiaohao Guo for assistance harvesting samples; John Zdenek and Tommy Clark for assistance with plant care; and Bader Arouisse, Ella Katz, Daniel Kliebenstein, Arthur Korte, Baptiste Mayjonade, and members of the Bergelson Lab for sharing data and/or helpful feedback. Amélie Vergnol was a student in the *Magistère de Génétique* Graduate Program at Université de Paris when the work was conducted.

