## Supplementary Note, Methods, Tables and Figures for "Genome-wide association mapping within a single *Arabidopsis thaliana* population reveals a richer genetic architecture for defensive metabolite diversity"

#### (a) Glucosinolate extraction methods.

Traditional approaches to GSL extraction are time-intensive. To facilitate high-throughput phenotyping, we explored modifications of the approach of [1], who demonstrated that extraction in 80% methanol is sufficient to inhibit GSL-degrading myrosinase activity even without lyophilization or boiling. Further, they showed that traditional methods may introduce variability into GSL profiles, including biased loss of some compounds.

Specifically, we quantified GSL profiles following extraction in 80% methanol while varying two steps: leaf samples were frozen beforehand or added to the methanol solution immediately after collection, and leaf samples were immediately homogenized while submerged in the methanol solution or left intact. Samples were subsequently incubated 48h in the dark at room temperature. Extractions were conducted on small rosette leaves (~200mg) in 1.5 mL polypropylene tubes from two representative TOU-A accessions with similar GSL profiles (N = 2 replicates per method per accession x 2 accessions).

Results of each approach were comparable (Supplementary Note, Figure 1), except GSLs were reduced in previously-frozen leaves that were not ground to initiate the extraction. This is likely due to activity of myrosinase as cells thawed before the methanol solution fully diffused through the tissue. When profiling 294 TOU-A accessions for GWAS, we therefore extracted GSLs from intact rosettes placed directly in 80% methanol without tissue homogenization.

#### (b) Glucosinolate quantification methods.

We implemented a simple approach that does not require delineating specific peak boundaries to quantify GSLs, with the goal of achieving high sensitivity for low-abundance GSLs (i.e., peaks that are less pronounced relative to background noise). Specifically, for a given m/z and retention time range, this approach infers the baseline value as the i-th (user specified) percentile value among all observed intensity values, and sums all distances from the baseline to the recorded intensity for all points in the upper j-th (user-specified) percentile. This is akin to integrating the area under a curve by summing up the lengths of lines that densely fill the space underneath; integrated peak areas can easily be adjusted post-hoc for the frequency of intensity observations per second if desired. Our pipeline produced plots of each curve (Supplementary Note, Figure 2) for every sample, which we inspected to determine optimal values of i and j for each molecule. The approach is straightforward to implement for a metabolite across thousands of samples in a few minutes, and scripts are provided as described in the Data Availability section of the main text. Because our approach produces non-zero estimates even in the absence of true peaks, “blank” samples can be inspected to determine expected background values to subtract from observed values; in our experience, however, background noise values were minimal even compared to minor peaks in our samples.

We compared our approach to peak detection and integration using the CentWave method implemented in the findChromPeaks function of *xcms* [2]. We optimized CentWave parameters (peakwidth, snthresh, prefilter) for each molecule by manually inspecting results across a range of values. We found that while both methods produced nearly identical results for the GSLs with the highest intensity peaks, our customized approach had higher sensitivity for lower-intensity peaks (Supplementary Note, Figure 3).

This performance of this method was supported by results for 4hIM, a molecule produced as a precursor to 4mOIM. Given that 4moIM was detected abundantly in every accession, we expected to find non-zero amounts of its 4hIM in every accession as well. This was the case using our customized approach, which found an approximately normal distribution of 4hIM abundances across samples; by contrast, the peak-picking approach found no peaks for 4hIM in the majority of samples (Supplementary Note, Figure 3).

Given its high sensitivity for low-abundance peaks and strong agreement with peak-picking methods for high-abundance peaks, we used our customized integration approach in R for high-throughput quantification of peaks across all TOU-A samples in our experiment.

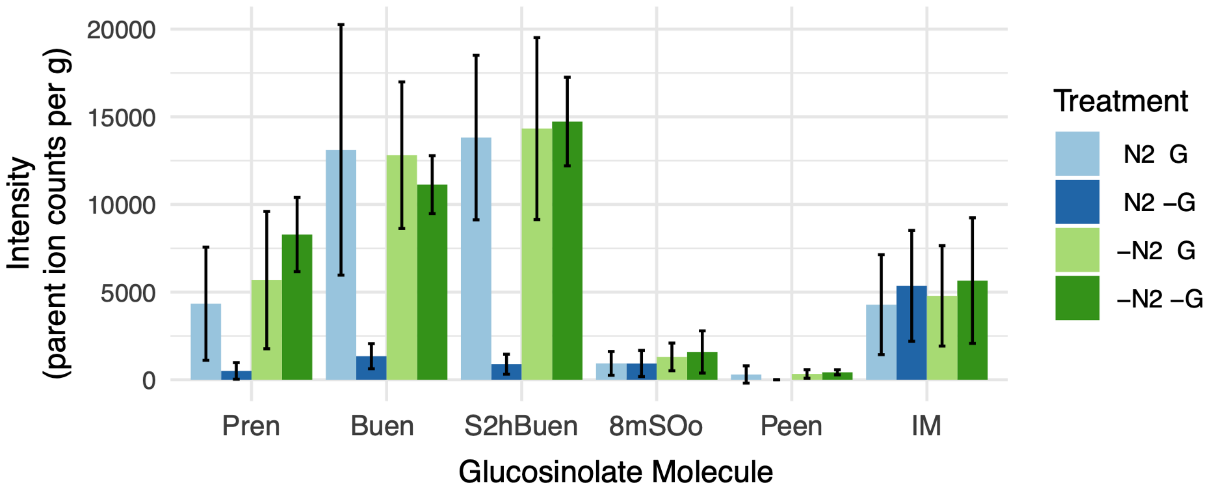


**Supplementary Note, Figure 1. Comparison of extraction methods for quantification of GSLs in TOU-A plants.**

Four samples per treatment were extracted in 80% methanol as described above. Treatment abbreviations: N2/-N2: with and without freezing tissue in liquid nitrogen before extraction; G/-G: with and without grinding sample to initiate extraction. -N2 -G corresponds to the extraction method used for our full study.

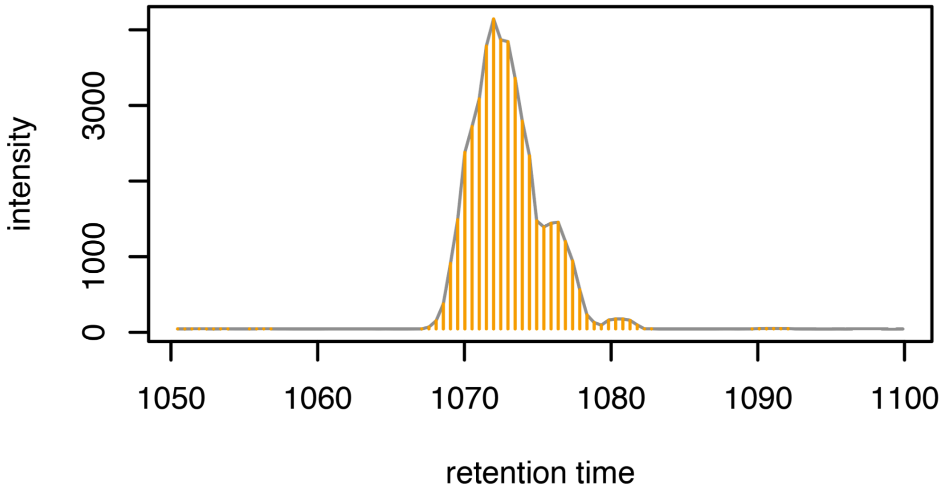


**Supplementary Note, Figure 2. A chromatogram peak from HPLC-MS/MS analysis of GSLs in a TOU-A sample, annotated to show the integrated peak area using our customized approach.**

Each orange line indicates the distance from the observed intensity to the inferred baseline, and peak area is determined as the sum of these distances. Points in the upper j-th (user-specified) percentile of intensity values within the retention time window were used for integration. Some lines are clearly outside the peak, but their contribution to total area is negligible, such that a wide retention time window and liberal value of j typically results in fully integrated GSL peaks across a range of samples, even when samples display some variation in retention time and peak shape.

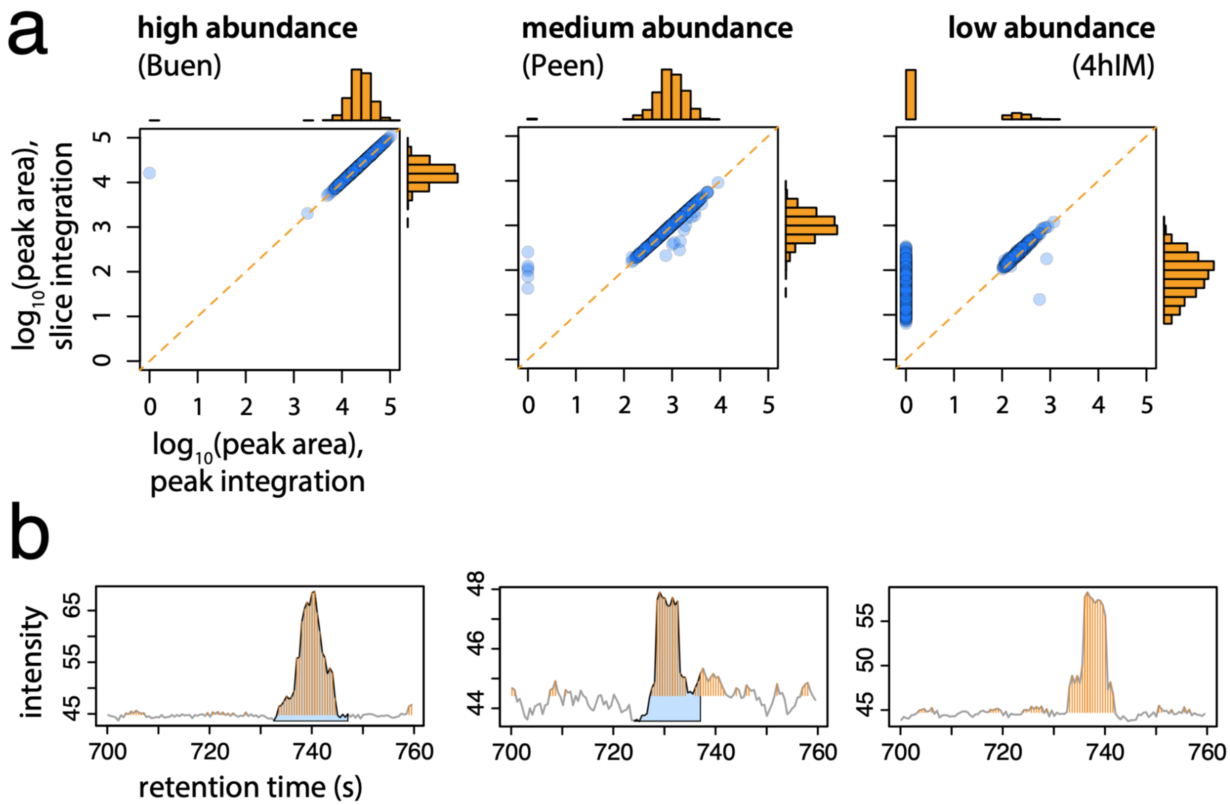


**Supplementary Note, Figure 3. Comparison of integration methods for glucosinolate quantification in TOU-A plants (N > 1,100 samples).**

**(a)** Inferred abundance (parent ion counts) using the customized slice integration method and the CentWave peak picking algorithm in *xcms* shown for glucosinolate molecules with high (left), medium (center), and low (right) abundance.
**(b)** Slices (orange) and defined peaks (blue) shown for an instance in which both methods were in close agreement (left) and for two representative instances of the main sources of disagreement (center and right). These examples were taken from the analysis of 4hIM, one of the lowest-intensity peaks for TOU-A. The peak picking method occasionally inferred a higher abundance when it identified a lower baseline intensity against a noisy background (center), or failed to detect a peak (right).

In panels (a) and (b), units for the y-axis are parent ion counts measured for the focal molecule.

### Supplementary Methods

**(a) SNP comparisons among TOU-A and 1001G accessions**

To compare measures of diversity between the two mapping panels, SNPs shared between the two mapping populations were analyzed. Various SNP sets were generated as described in the Results of the Main Text using bcftools version 1.11, and only included biallelic TOU-A SNPs with a “PASS” status indicated from SNP calling by [3] (which excludes SNPs with a ‘str10’ flag, in which > 90% of reads mapped to one of the strand). SNPs were compared to those in the 1001 Genomes panel.

**(b) Allele frequency spectrum**

To compare distributions of allele frequencies across 1001 genomes and TOU-A, variants were filtered using bcftools v. 1.11 as described above. We utilized the 1001G genotypes available on the project’s webserver (https://1001genomes.org/data/GMI-MPI/releases/v3.1/). For 1001G, a random subset of 195 European accessions was taken to match the sample size of TOU-A genomes in our analysis, and subsequently limited to biallelic sites. For each mapping panel, sites in which 100 or more individuals were genotyped at a site were downsampled to 100 alleles randomly drawn without replacement from the total pool of genotyped alleles at that locus. A new minor allele frequency was then calculated. This was done to avoid any allele frequency binning bias by a population-dependent denominator, as well as biases in the number of genotyped individuals between panels at particular loci. The density of these downsampled minor allele frequencies was plotted as the proportion of SNPs present in 1% minor allele frequency bins, from 1 to 50.

Tajima’s D was also calculated using vcftools v. 0.1.16 in each panel using the above downsampled minor allele frequencies in windows of 50,000 bp.

**(c) Population structure**

For 1001G, minor alleles from genomic chromosomes were filtered at a frequency of 0.03. For the resulting 2,620,695 SNPs, a haplotype filter of 0.8 $r^2$ was applied in PLINK v. 1.9.0, resulting in 921,495 markers in relative linkage equilibrium. TOU-A was similarly filtered, with 1,282,049 SNPs passing the 0.03 maf filter and 239,309 markers passing the LD threshold.

These marker sets were then used in ADMIXTURE v. 1.3.0 with K values ranging from 1 to 15. The estimation failed to converge for larger values of K (K = 7-15) in the 1001 genomes marker set. These higher values of K were discarded in further analyses for both 1001G and TOUA. However, for both TOUA and 1001 genomes, error values associated with cross-validation (produced with default settings in ADMIXTURE) decreased with increasing K.

In 100KB windows around each locus harboring a significant GSL association in our GWAS within TOU-A, the strength of population structure was inferred by taking the variance of the five estimated ancestral allele frequencies (i.e., within-group allele frequencies for K=5) for each site, and comparing these values with the minor allele frequencies per site. Loess fits of each distribution were estimated separately for TOU-A and 1001G and include standard error. Windows of 30KB and 200KB and different values of K showed similar patterns (results not shown).

**(d) Glucosinolate chemotypic variation as a function of geographic distance**

Using accessions from Katz et al. [4], the geographic distance between every accession pair was calculated using the Haversine formula (R package geosphere v. 1.5-10) applied to collection locality latitude and longitude coordinates. Using glucosinolate chemotype classifications assigned by Katz et al., ‘mismatches’ were determined by whether two accessions shared the same chemotype classification. The proportion of mismatches was binned and plotted, and fits of each distribution were estimated using loess.

### Supplementary Tables

**Table S1. Glucosinolate molecules quantified in the TOU-A population with corresponding mass-to-charge ratio and retention time ranges.**

The numeric "id" for each molecule refers to the numbering system proposed by [5] and continued by [6,7].

| id | name | abbreviation | alt. abbreviation | m/z | ret. time (s) |
|---|---|---|---|---|---|
| 73 | 3-methylsulfinylpropyl | 3mSOp | 3MSP | 421.5-422.5 | 280-400 |
| 24R | R2-hydroxy-3-butenyl | R2hBuen | 2H3B | 387.5-388.5 | 380-490 |
| 107 | 2-propenyl | Pren | 2P | 357.5-358.5 | 400-600 |
| 24S | S2-hydroxy-3-butenyl | S2hBuen | 2H3B | 387.5-388.5 | 490-600 |
| 64 | 4-methylsulfinylbutyl | 4mSOb | 4MSB | 435.5-436.5 | 500-600 |
| 72 | 5-methylsulfinylpentyl | 5mSOp | 5MSP | 449.6-450.6 | 580-650 |
| 38 | 2-hydroxy-4-pentenyl | 2hPeen | 2H4P | 401.6-402.6 | 595-680 |
| 12 | 3-butenyl | Buen | 3B | 371.5-372.5 | 600-700 |
| 67 | 6-methylsulfinylhexyl | 6mSOh | 6MSH | 463.6-464.6 | 630-710 |
| 28 | 4-hydroxy-indol-3-ylmethyl | 4hIM | 4HI3M | 462.5-463.5 | 700-760 |
| 66 | 7-methylsulfinylheptyl | 7mSOh | 7MSH | 477.6-478.6 | 700-800 |
| 101 | 4-pentenyl | Peen | 4P | 385.6-386.6 | 720-820 |
| 69 | 8-methylsulfinyloctyl | 8mSOo | 8MSO | 491.6-493.1 | 790-850 |
| 43 | indol-3-ylmethyl | IM | I3M | 446.6-447.6 | 800-900 |
| 48 | 4-methoxy-indol-3-ylmethyl | 4moIM | 4MOI3M | 476.6-477.6 | 850-950 |
| 47 | 1-methoxy-indol-3-ylmethyl | 1moIM | 1MOI3M | 476.6-477.6 | 950-1050 |
| 87 | 7-methylthioheptyl | 7mSh | 7MTH | 461.6-462.6 | 1050-1100 |
| 92 | 8-methylthiooctyl | 8mSo | 8MTO | 475.6-477.1 | 1070-1150 |

### Supplementary Figures

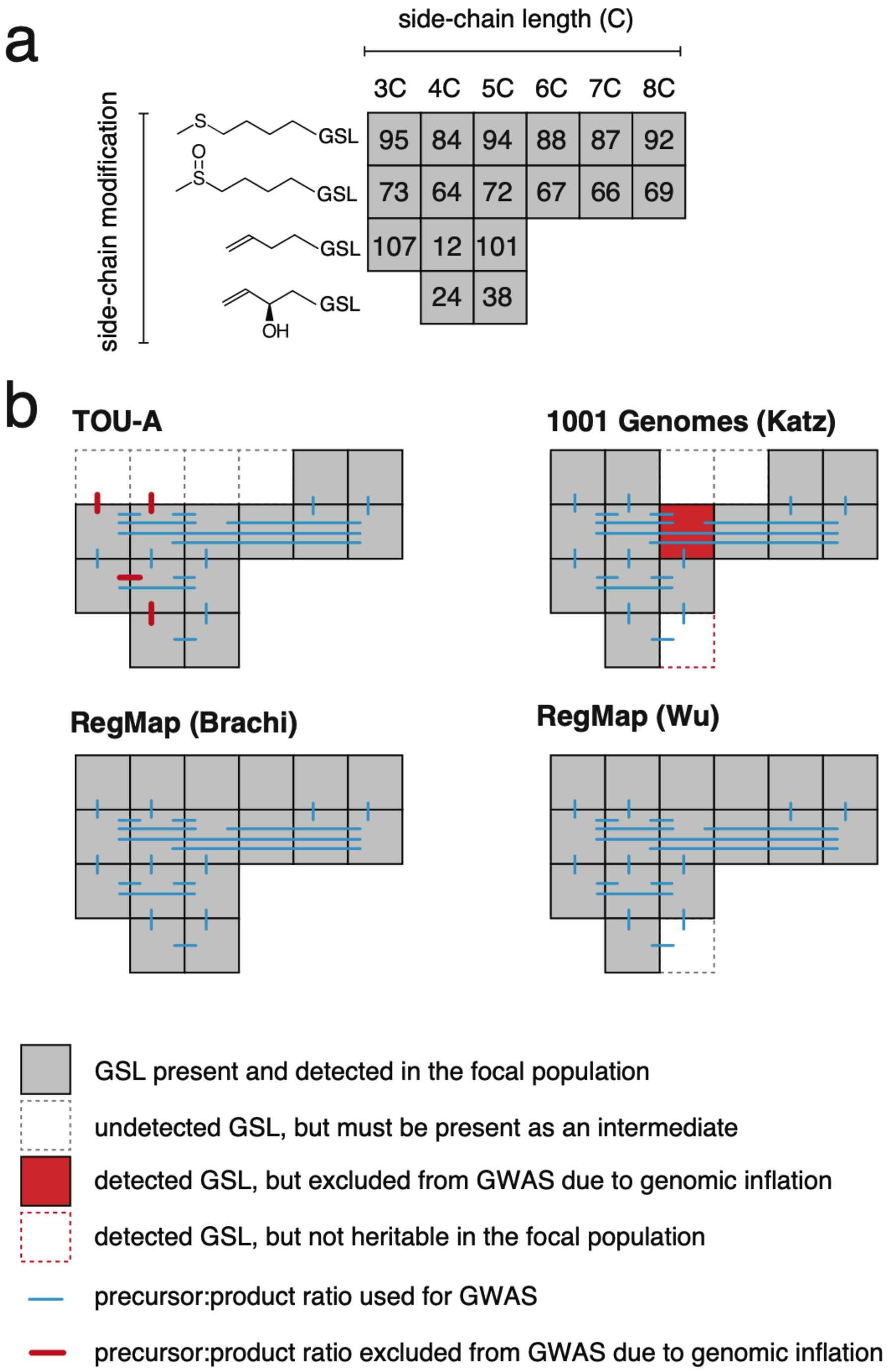

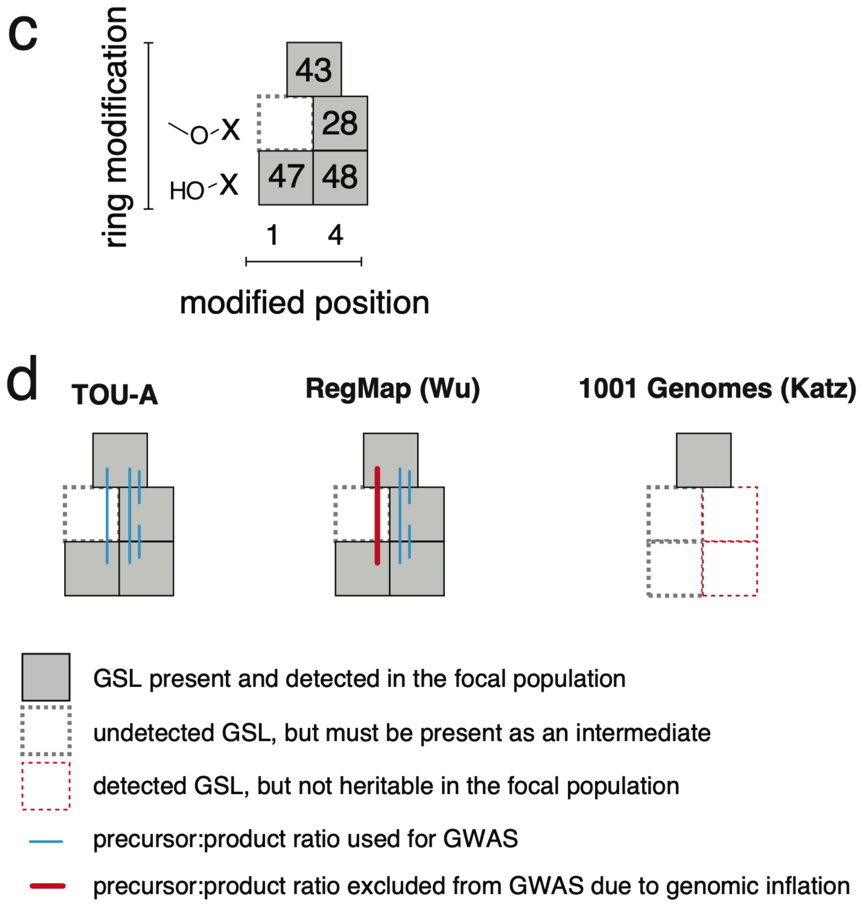


**Figure S1. Glucosinolate molecules and precursor:product ratios quantified in each population and used in GWAS.**

**(a)** Aliphatic GSL molecules quantified across studies in the TOU-A, RegMap, and 1001G populations. Boxes represent GSL molecules, with their numeric IDs from Table S1 (taken from [5]). Note that two molecules that are the product of an active AOP3 enzyme (IDs 25 and 42) were not produced in TOU-A plants and are not shown in the figure.
**(b)** Summary of aliphatic molecules detected in each population.
**(c)** Indolic GSL molecules, represented as in panel a.
**(d)** Summary of indolic molecules detected in each population.

Note that vertical lines represent direct precursor:product relationships among the indicated molecules. Horizontal lines represent indirect precursor:product relationships, arising from direct precursor:product relationships upstream within the MAM-catalyzed side-chain elongation cycle that acts on GSL precursor molecules.

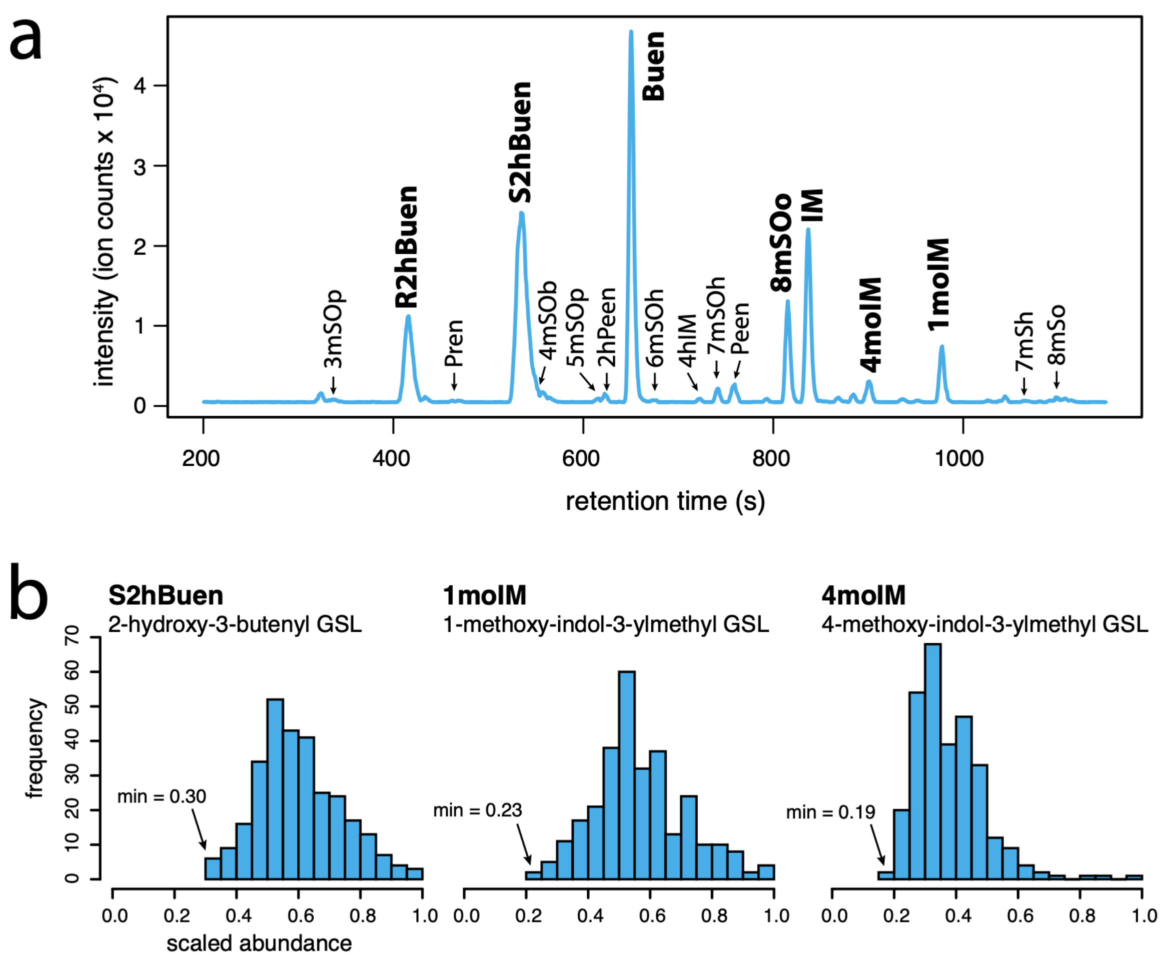


**Figure S2. Glucosinolate profiles in the TOU-A population indicate an absence of large effect loss-of-function variants within the glucosinolate biosynthetic pathway.**

**(a)** Extracted ion chromatogram (EIC) from HPLC-MS/MS quantification of GSLs in a sample from the TOU-A population. The plot shows intensity values per g leaf tissue for the full range of parent ion mass-to-charge (m/z) ratios that was monitored (see Methods). Although some minor peaks are overlapping, note that individual peaks were extracted separately using narrow m/z ranges.
**(b)** Distributions of genetic values (BLUPs) across TOU-A accessions for three molecules that are terminal products of GSL biosynthesis in *Arabidopsis* (excluding very low-abundance derivatives of these molecules, if any). Values are scaled by the maximum value observed within the population. S2hBuen is the terminal product in short-chain aliphatic GSL biosynthesis from Methionine and requires functional GS-OX(n), MAM1, AOP2, and GS-OH enzymes. 1moIM and 4moIM are alternative terminal products of indole GSL biosynthesis from Tryptophan and require one or more functional IGMT(n) and CYP81F(n) enzymes. The "(n)" suffix refers to enzymes with multiple, sequentially numbered paralogs.

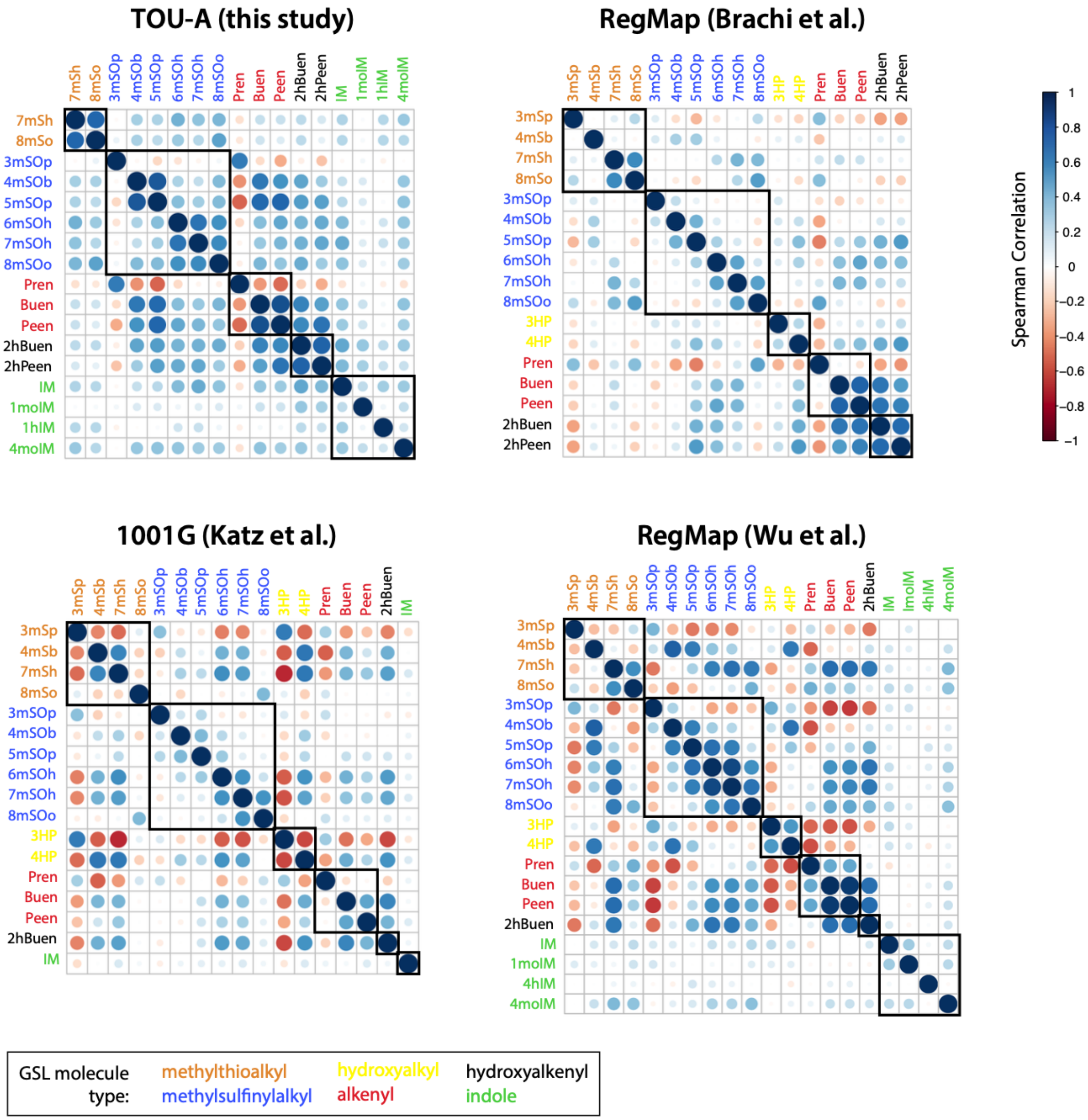


**Figure S3. Genetic correlations within the glucosinolate biosynthetic pathway.**

Spearman correlation coefficients among the model-fitted genotypic effects (BLUPs or means per accession; see Methods) for each GSL analyzed in this study. Names of GSLs with the same structural type side-chain modification are color-coded, and corresponding correlation coefficients for these related molecules are grouped by black boxes within the plot. The mapping panel and source of the data are indicated above each plot.

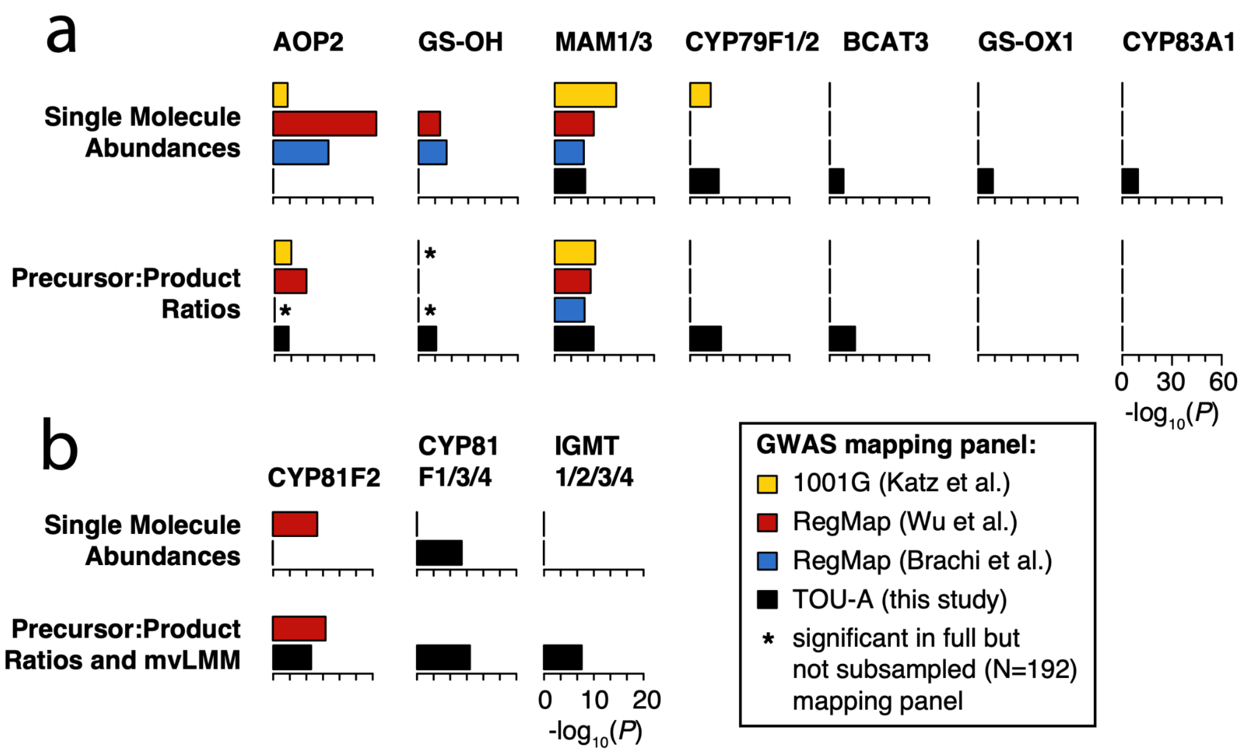


**Figure S4. Significant associations across mapping panels at five biosynthetic loci for aliphatic and indolic glucosinolates.**

**(a)** Among all SNPs assigned to each aliphatic GSL biosynthetic locus with significant associations in any population (see Methods), the top *P*-value was determined across GWAS for all molecular abundances (upper row) or precursor:product ratios (lower row). Datasets for the RegMap and 1001G mapping panels were randomly subsampled to 192 accessions, the sample size used for the TOU-A population, prior to conducting GWAS. Significant associations were determined using a Bonferroni correction accounting for both the number of SNPs investigated and the number of individual GWAS per trait type (i.e., the number of molecules for which individual GWAS were conducted).
**(b)** Plots were produced as in panel a, but for SNPs assigned to indolic GSL biosynthetic loci. Note that the mvLMM multi-trait GWAS model, plotted together with GWAS for individual precursor:product ratios, could not be conducted for the Wu et al. dataset due to algorithmic termination errors. Indolic GSLs from Katz et al. were excluded due to low heritability.

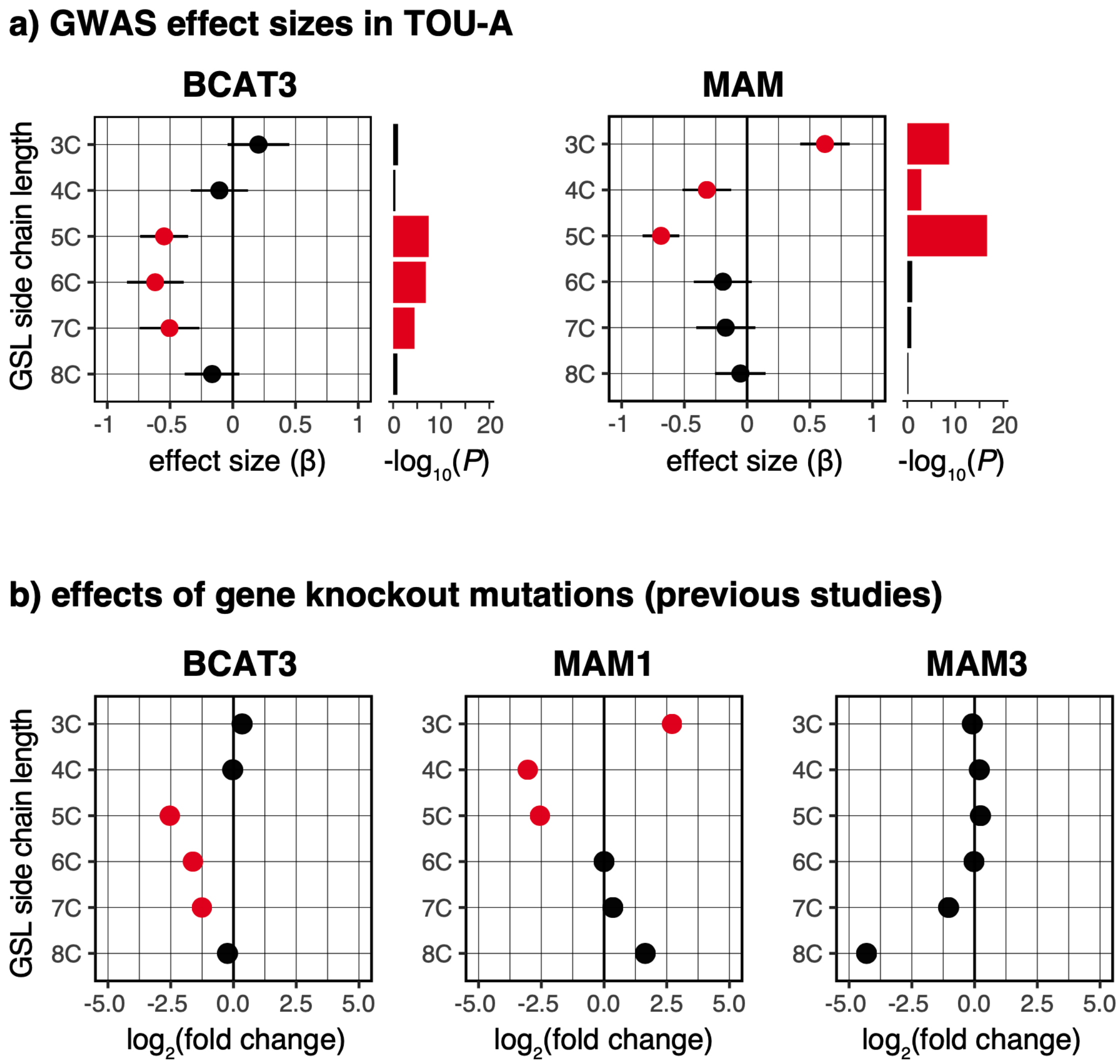


**Figure S5. BCAT3 underlies an additional component of glucosinolate chain length variation in TOU-A, consistent with phenotypes in gene knockout mutants.**

**(a)** Effect sizes (units = phenotypic standard deviations, ± 95% CI) and *P*-values for the leading SNP in individual GWAS models for the abundance of each methylsulfinylalkyl glucosinolate in TOU-A. The leading SNP was defined as the SNP with the strongest study-wide *P*-value at the locus, not on a per-trait basis. Data points for the three molecules with the largest effect sizes in each plot are colored red.
**(b)** Fold-change in the abundance of the molecules from (a) in knockout mutant relative to wild-type *Arabidopsis* (Col-0 accession), adapted from previously published data [8] to indicate the concordance between patterns in panels (a) and (b). Points are colored as in (a) to highlight concordant patterns.

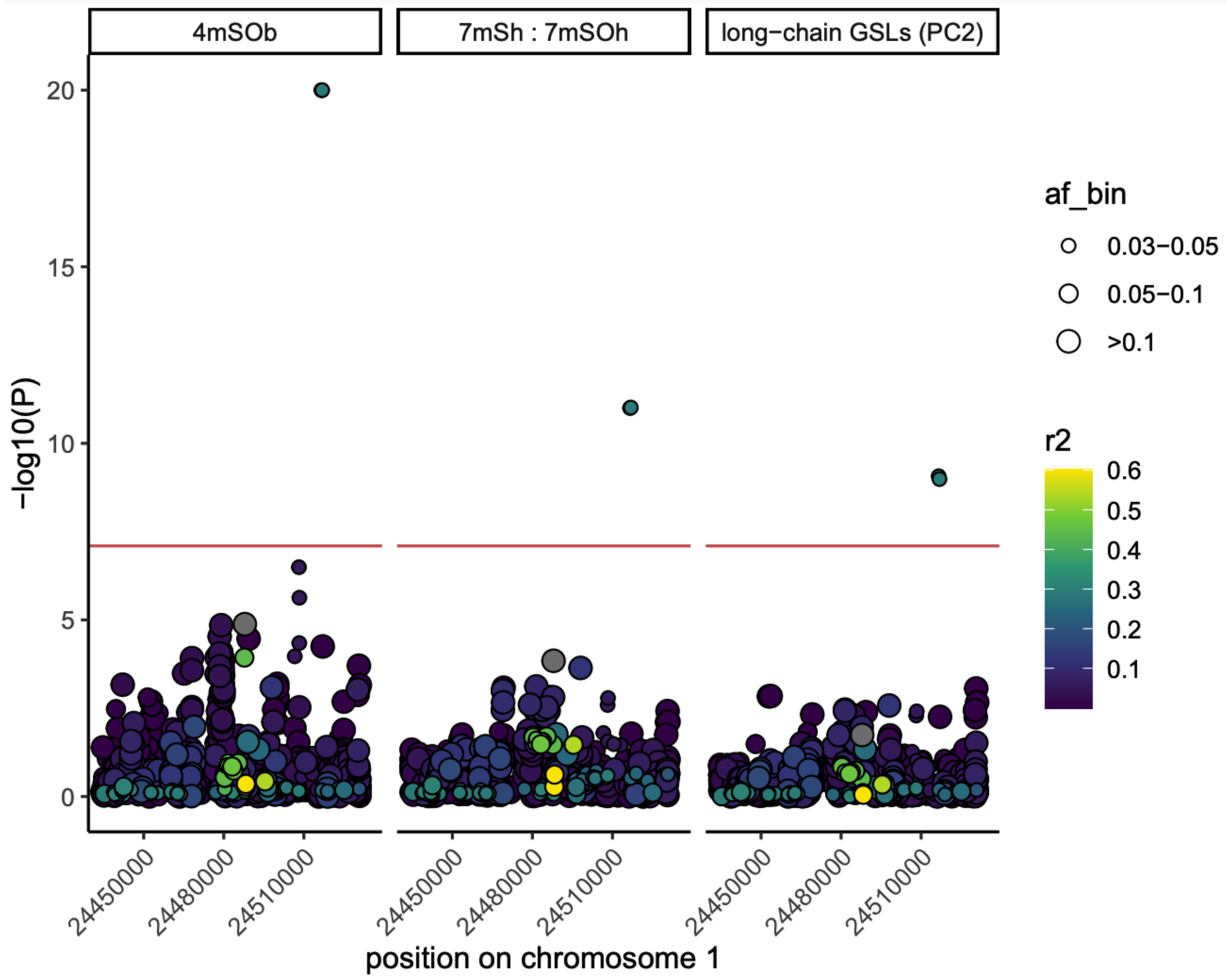


**Figure S6. Associations at a minor variant near GS-OX1 in the TOU-A population.**

*P*-values per SNP are shown for three GWAS, centered on the SNP (colored gray) that consistently exhibits the strongest associations among common variants (maf > 0.05, colored gray) within 30 kb of GS-OX1. The red line indicates the Bonferroni-corrected significance threshold. Colors for other SNPs indicate LD ($r^2$) with this SNP, and point sizes indicate minor allele frequency bins ("af_bin"). Note that a slightly lower frequency allele (maf = 0.03) in LD with this SNP exhibits the strongest associations, and this variant reaches statistical significance for two traits expected to be impacted by GS-OX enzymes: the precursor:product ratio for 7mSo:7mSOo GSLs, and a principal component capturing divergent effects on xmSo (7mSo, 8mSo) vs. xmSOo (6mSOo, 7mSOo, 8mSOo) GSLs in TOU-A.

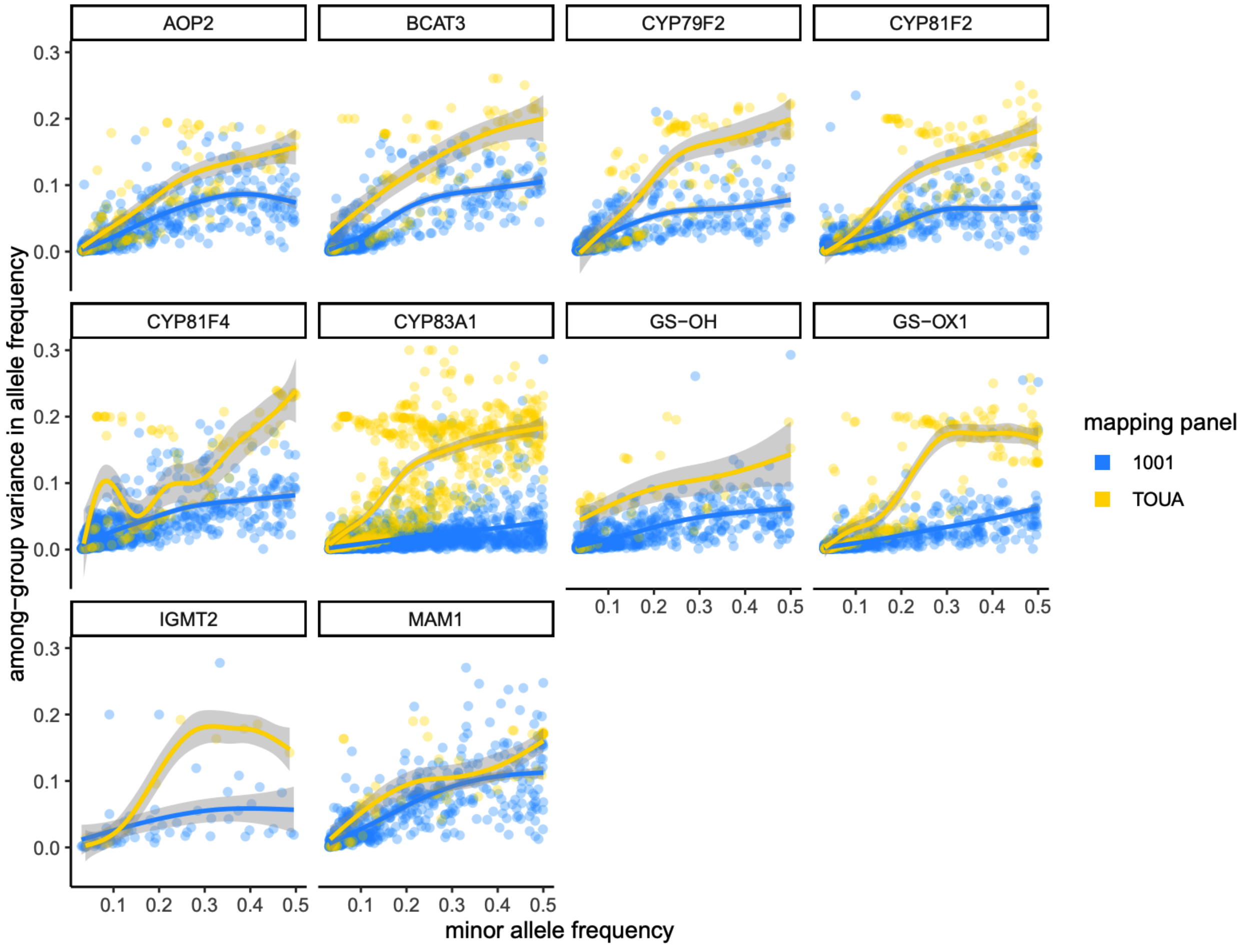


**Figure S7. Divergence in allele frequency among subpopulation groups is not exaggerated across Europe relative to divergence within TOU-A.**

Variance in allele frequencies among subpopulation groups inferred using ADMIXTURE (k = 5), plotted as a function of the minor allele frequency, for SNPs within 100kb of putatively causal genes at ten loci significantly associated with glucosinolate variation in TOU-A. Note that values are not elevated (and are typically reduced) in the European 1001G panel relative to the local TOU-A population; thus, SNPs at these loci do not appear to track population structure more strongly over a broader geographic scale.
